## Supplementary figures and images for "CircRNA_012164/MicroRNA-9-5p axis mediates cardiac fibrosis in diabetic cardiomyopathy"

### Supplemental Figure 1

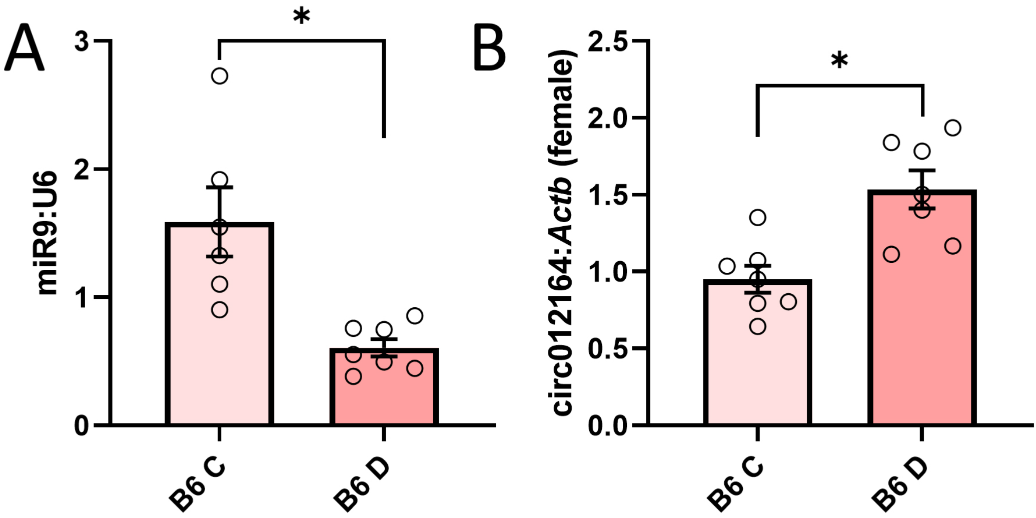

### Supplemental Figure 2

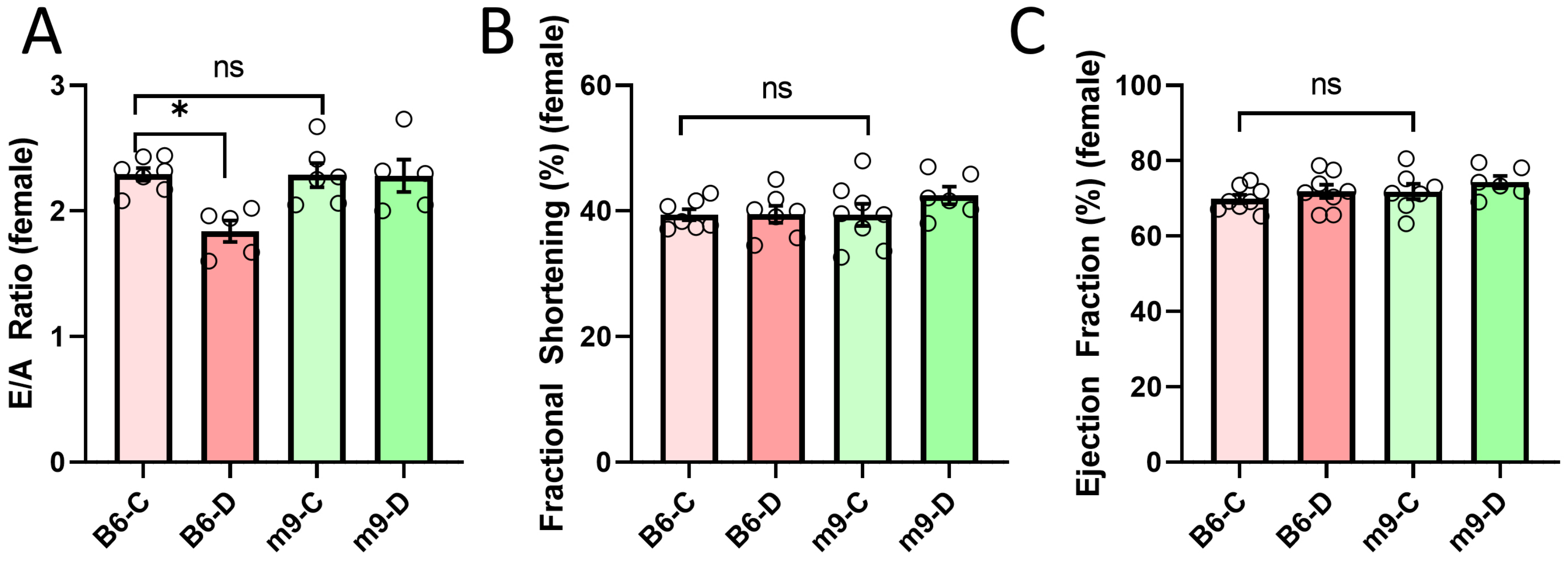
