## Supplemental Table 1 for "CircRNA_012164/MicroRNA-9-5p axis mediates cardiac fibrosis in diabetic cardiomyopathy"

| siRNA | Sense 5'-3' | Antisense 5'-3' |
|---|---|---|
| **Anti-circRNA_012164_1** | UGCCAAGCCCAAGG UCACCAAGCUU | GCUUGGUGACCUUG GGCUUGGCAUU |
| **Anti-circRNA_012164_2** | GCCCAAGGUCACCA AGCCCAAGAUU | CUUGGGCUUGGUGA CCUUGGGCUU |
